# Different DNA repair pathways are required following excision and integration of the DNA cut & paste transposon *piggyBat* in *Saccharomyces cerevisiae*

**DOI:** 10.1101/015289

**Authors:** Weifeng She, Courtney Busch Cambouris, Nancy L. Craig

**Affiliations:** Howard Hughes Medical Institute, Department of Molecular Biology & Genetics, Johns Hopkins University School of Medicine, Baltimore, MD 21205

**Author notes:** Corresponding author: Nancy L. Craig, Howard Hughes Medical Institute, Department of Molecular Biology & Genetics, Johns Hopkins School of Medicine, 502 PCTB, 725 N. Wolfe Street, Baltimore, MD 21205, voice =, email =, fax =.

## Abstract

The movement of transposable elements from place to place in a genome requires both element-encoded and host-encoded factors. In DNA cut & paste transposition, the element-encoded transposase performs the DNA breakage and joining reactions that excise the element from the donor site and integrate it into the new insertion site. Host factors can influence many aspects of transposition. Notably, host DNA repair factors mediate the regeneration of intact duplex DNA necessary after transposase action by repairing the double strand break in the broken donor backbone, from which the transposon has excised, and repairing the single strand gaps that flank the newly inserted transposon. We have exploited the ability of the mammalian transposon *piggyBat*, a member of the *piggyBac* superfamily, to transpose in *Saccharomyces cerevisiae* and used the yeast single gene deletion collection to screen for genes encoding host factors involved in *piggyBat* transposition. *piggyBac* transposition is distinguished by the fact that *piggyBac* elements insert into TTAA target sites and also that the donor backbone is restored to its pre-transposon sequence after transposon excision, that is, excision is precise. We have found that repair of the broken donor backbone requires the non-homologous end-joining repair pathway (NHEJ). By contrast, NHEJ is not required for DNA repair at the new insertion site. Thus multiple DNA repair pathways are required for *piggyBac* transposition.

## INTRODUCTION

Transposable elements are mobile DNA segments that have the ability to move from one position to another in a genome (Craig *et al*. 2002). They have significant impact on chromosome structure, function, and evolution in virtually all organisms and thus mediate multiple fundamental processes in biology. We are particularly interested in DNA cut-and-paste transposition in which an element-encoded transposase binds specifically to the transposon ends and mediates the DNA breakage and joining reactions that excise the element from the donor site and integrate it into a new insertion site (Peters and Craig 2001a; Peters and Craig 2001b). Studies in many systems have revealed that host factors can influence many aspects of transposition. Host factors can modulate the level of transposase, facilitate the assembly of the protein-DNA complexes in which transposition occurs by promoting DNA bending, channel insertions to particular target sites by interacting with the transposase and the target DNA, or facilitate the disassembly of highly stable post-transposition complexes (Wardle *et al*. 2009). Host DNA repair factors also play key roles in transposition. DNA repair reactions at the donor site and at the new insertion site are required to complete transposition by regenerating intact duplex DNA after transposase-mediated DNA breakage and joining. Excision of the transposon from the donor site by DNA double-strand breaks at the transposon ends leaves a double-strand gap in the donor DNA which must be repaired. Furthermore, the newly inserted transposon is flanked by single-strand gaps which also must be repaired. Repair of these gaps generates the target site duplications that flank the newly inserted transposon.

Here we exploit the genetic tractability of the yeast *Saccharomyces cerevisiae* and use the single gene deletion collection to screen for host factors involved in the excision and donor site repair of a mammalian *piggyBac* cut & paste transposon that is also active in yeast. As described in Results, our screen revealed that the NHEJ pathway is required for ligation of the gapped donor backbone.

*piggyBat*, a member of the *piggyBac* superfamily (Fraser *et al*. 1996) found in the little brown bat, *Myotis lucifigus* (Ray *et al*. 2008), was the first active mammalian DNA cut & paste transposable element to be identified (Mitra *et al*. 2013). We showed directly that *piggyBat* is an active transposon by assay of both transposon excision and integration in bat and human cells and in *S. cerevisiae*. As previously established with insect *piggyBac (Mitra et al. 2008)*, *piggyBat* inserts specifically at TTAA target sites and excises precisely, that is the donor site is restored to its pre-insertion TTAA sequence.

The transposase-mediated DNA breakage and joining reactions that underlie *piggyBac* transposition was established by *in vitro* analysis of insect *piggyBac* transposition and involves four distinct steps (Figure 1) (Mitra *et al*. 2008). Because *piggyBac* inserts specifically into TTAA target sites, *piggyBac* in a donor site is flanked by TTAA direct repeats. Transposition is initiated by transposase nicking at the 3’ ends of the transposon, followed by the attack of the newly exposed 3’OHs on the complementary donor strand four nt into the flanking donor DNA, forming TTAA hairpins on both transposon ends and releasing the transposon from the donor DNA site. The 5’ ends of the donor gap also have TTAA overhangs. Rejoining of these overhangs can restore the donor site to its pre-transposon state, resulting in precise excision.

**Fig 1.**
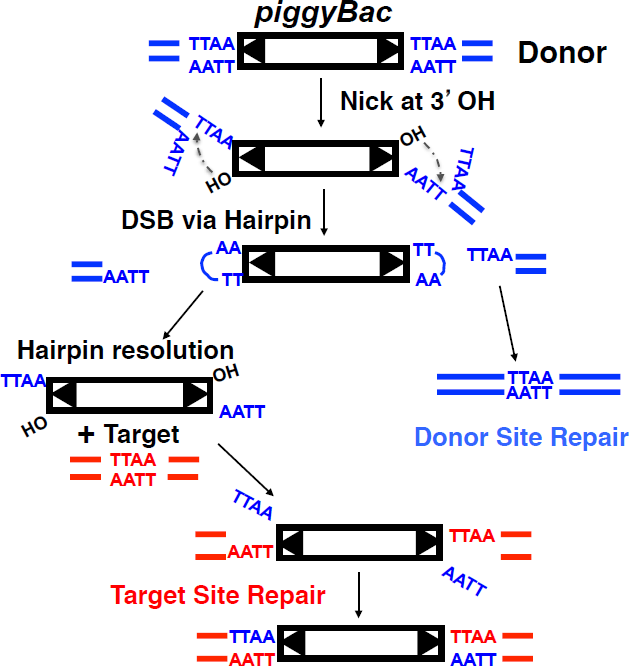
*piggyBac* transposition pathway. *piggyBac* transposition is initiated by nicks at the 3’ transposon ends. The exposed 3’OHs then attack the complementary strand 4 nt inside the flanking donor DNA to form the hairpins on the transposon ends. Once the transposon is released from the donor site, the double strand break in the donor backbone is precisely repaired to the pre-insertion TTAA sequence. The hairpins on the transposon ends are nicked at the 3’ transposon ends to expose the 3’OHs. The 3’OH transposon ends attack the TTAA target sequence at staggered positions, forming covalent links between the 3’ ends of the transposon and the 5’ ends of the target site. The single strand gap between the 3’ ends of the target DNA and the 5’ ends of the transposon are repaired to generate the four bp TTAA target sequence duplication.

The TTAA hairpins on the excised *piggyBac* transposon ends are resolved by the transposase to form four nt TTAA overhangs at the 5’ ends of the transposon. The released 3’OH transposon ends then attack the target TTAA sequence at staggered positions, covalently joining the transposon to the target DNA such that the transposon is flanked by single strand gaps. Repair of these gaps could occur by simple ligation of the TTAAs on the 5’ transposon ends to the target DNA or by a gap filling mechanism to form the target site duplications.

What host factors mediate these repair reactions at the gapped donor site and the insertion site? Studies of eukaryotic transposons have revealed that both non-homologous end-joining (NHEJ) and homology-dependent repair can be used to repair the donor site gaps (Mcvey *et al*. 2004; Yant and Kay 2003). The relative use of particular repair pathways is influenced by many factors including the nature of the donor stand breaks which differ for different superfamilies of elements, the host organism, the stage of the cell cycle or whether transposition is examined in the germline.

During NHEJ repair of the donor backbones, the broken DNA ends are often rejoined with limited processing such that “footprints” of the element and its flanking target site duplication usually remain following excision. Thus most transposon donor sites are not restored precisely to their pre-transposon sequence. Alternatively the gap may be repaired by homology-dependent repair in which repair is templated by a sister chromatid such that a copy of the transposon is regenerated at the donor site or, if the homolog lacks a copy of the transposon, the gapped donor site will be repaired without copying a transposon, resulting in apparent precise excision.

We show here that rejoining of the gapped *piggyBat* donor backbone, which contains TTAA sequences on their 5’ ends, requires the NHEJ pathway in yeast. Simple ligation of these overhangs restores the donor site to its pre-transposon insertion state. We also show that repair of the gaps at the new insertion site does not require NHEJ and thus must use the non-NHEJ ligase Cdc9 ligase. Thus the regeneration of intact duplex DNA following *piggyBat* transposition requires multiple DNA repair activities despite the fact that both donor site and target site repair requires ligation of complementary overhanging TTAA sequences.

## METHODS

### Yeast strains

BY4727 *MATα his3Δ200, leu2Δ0, lys2Δ0, met15Δ0 trp1Δ63* (Brachmann *et al*. 1998) BY4741 *MATa his3*Δ*1 leu2*Δ*0 met15*Δ*0 ura3*Δ*0* (Brachmann *et al*. 1998) BY4741 *MATa his3*Δ*1 leu2*Δ*0 met15*Δ*0 ura3*Δ*0* Single gene deletion strains (Giaever *et al*. 2002)

### Construction of pWSY1-Excision

pWSY1-Excision is a pRS414-based plasmid containing the *piggyBat* transposase gene under the control of the GALS promoter and a mini-*piggyBat* transposon, which contains a NatMX resistance cassette, inserted within *URA3*, forming *ura3::* mini-*piggyBat-Nat*. First, the *TRP1* gene on the pRS414 *GALS piggyBat* transposase plasmid pGB50 described by Mitra et al (Mitra *et al*. 2013) was replaced with *URA3* by yeast homologous recombination in BY4727, using a PCR-generated *URA3* fragment flanked by 45 bp of the plasmid backbone, selecting for Ura^+^. The mini-*piggyBa*t-*Nat* was constructed by cloning a NATMX cassette bounded by HindIII and SphI sites amplified from pSG37 (Gangadharan *et al*. 2010) by PCR into a pUC plasmid containing the *piggyBat* ends L513 and R208 separated by a polylinker containing HindIII and SphI sites. The *mi*ni-*piggyBat-Nat* element was introduced into the first TTAA (bp 64) of *URA3* by homologous recombination using a *mi*ni-*piggyBat-Nat* fragment amplified from the pUC plasmid by PCR using primers complementary to *URA3*, selecting for resistance to clonNAT.

### *piggyBat* excision and donor site repair assay

A BY4727 pWS1-Excision colony freshly streaked on a SC + 2% glucose + 50 µg/ml clonNAT plate was inoculated into 5 ml of the same media and grown overnight at 30 °C. The cells were spun down, resuspended in 5 ml SC + 2% galactose + 50 µg/ml clonNAT and grown for 5 hours at 30°C. Cells were serially diluted and plated separately on YPD plates to determine the total number of cells and on SC - Ura + 2% glucose plates to determine the number of Ura^+^ cells in which excision occurred. The excision frequency is Ura^+^ cells/ total cells.

### The genome-wide screen

The yeast single gene deletion library in BY4741 was transformed *en masse* with the pWS1-Excision plasmid that carries a galactose-regulatable *piggyBat* transposase gene and *ura3::*mini-*piggyBat-Nat*, selecting for growth on SC + 2% glucose + 50 µg/ml clonNAT plates at 30°C. We performed three independent biological replicates starting from transformation of the library, obtaining at least 3 million transformants for each replicate. After growth for 48 hours, the transformants were recovered from the plates, concentrated, resuspended in 15% glycerol, and stored at −80°C.

To induce expression of the Gal-regulated *piggyBat* transposase, 25 OD_600_ of transformed cells were inoculated into 200 ml SC + 2% galactose + 50 µg/ml clonNAT and grown with shaking at 30°C for 5 hours. The cells were then recovered by centrifugation, washed and then resuspended.

To select cells in which *piggyBat* excision and repair of the *ura3::*mini-*piggyBat-Nat* donor site occurred, the induced transformants were grown-out in SC - Ura + 2% glucose. 25 OD_600_ of the galactose-induced cells were grown in 1 liter at 30°C for 24 hours, repeating this selection process twice more such that the cells grew for about 25 generations in selective media. Plating on YPD and SC - Ura + 2% glucose revealed that more than 90% of the selected cells were indeed Ura^+^.

### Barcode recovery and analysis

We recovered the deletion library UPTAG barcodes after transposase induction and after selection for excisants by PCR and quantitatively compared these pools by next generation sequencing. The primers included an 8-nucleotide multiplex index sequence that harbored the Illumina-sequencing primer sequence. We combined the six UPTAG samples in a single run on the Illumina Miseq platform and about 0.2 million reads were obtained for each sample. The Illumina sequencing reads were assigned to different samples using the 8-nucleotide multiplex index sequences from cycle 1 to cycle 8. The 20-mer sequences from cycle 27 to cycle 46 were matched with the updated UPTAG barcode sequences using Bowtie (Langmead and Salzberg 2012). To avoid dividing by zero, we added a pseudo-count of 1 to all reads before calculating the normalized fold changes. We then mapped the reads to the yeast genome (Smith *et al*. 2009). In Table 1, only gene deletions are listed for which there are more than 10 reads after transposase induction was present for each replicate. Among the deletion strains with log2 ratio >2.7, results from *Δynl296W*, *Δydl041W and Δynr005C* were omitted *because* their ORFs are dubious.

**Table 1.**
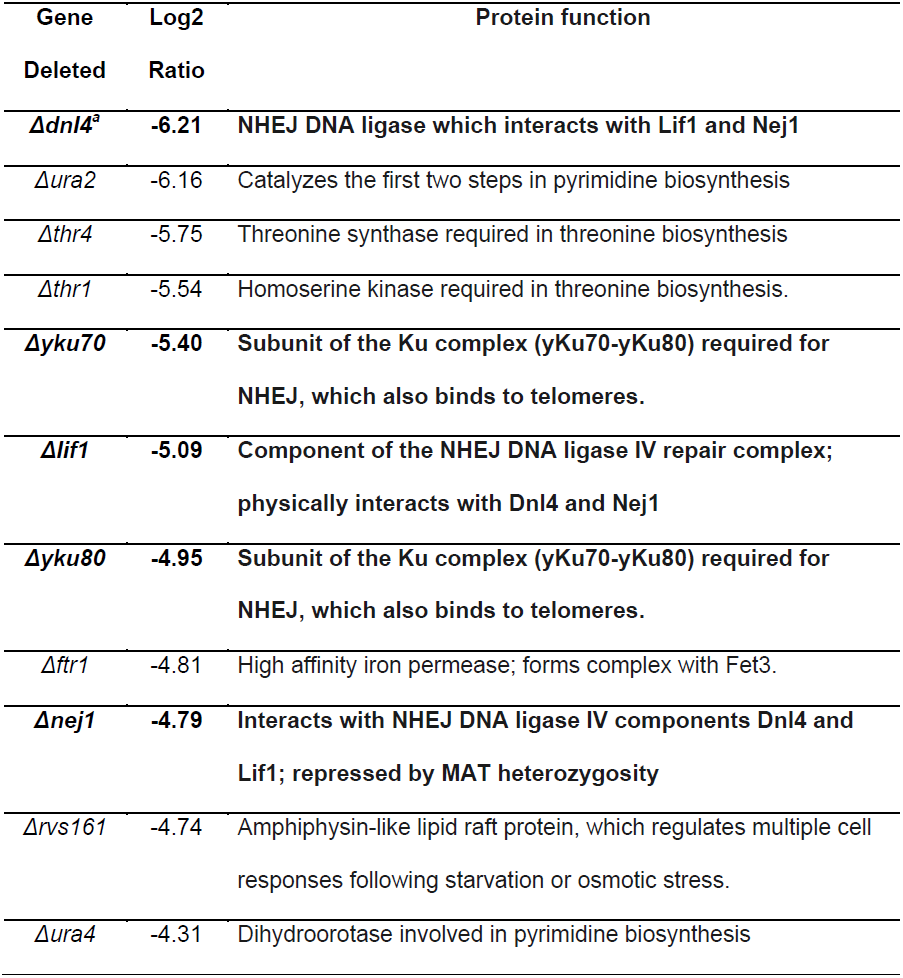
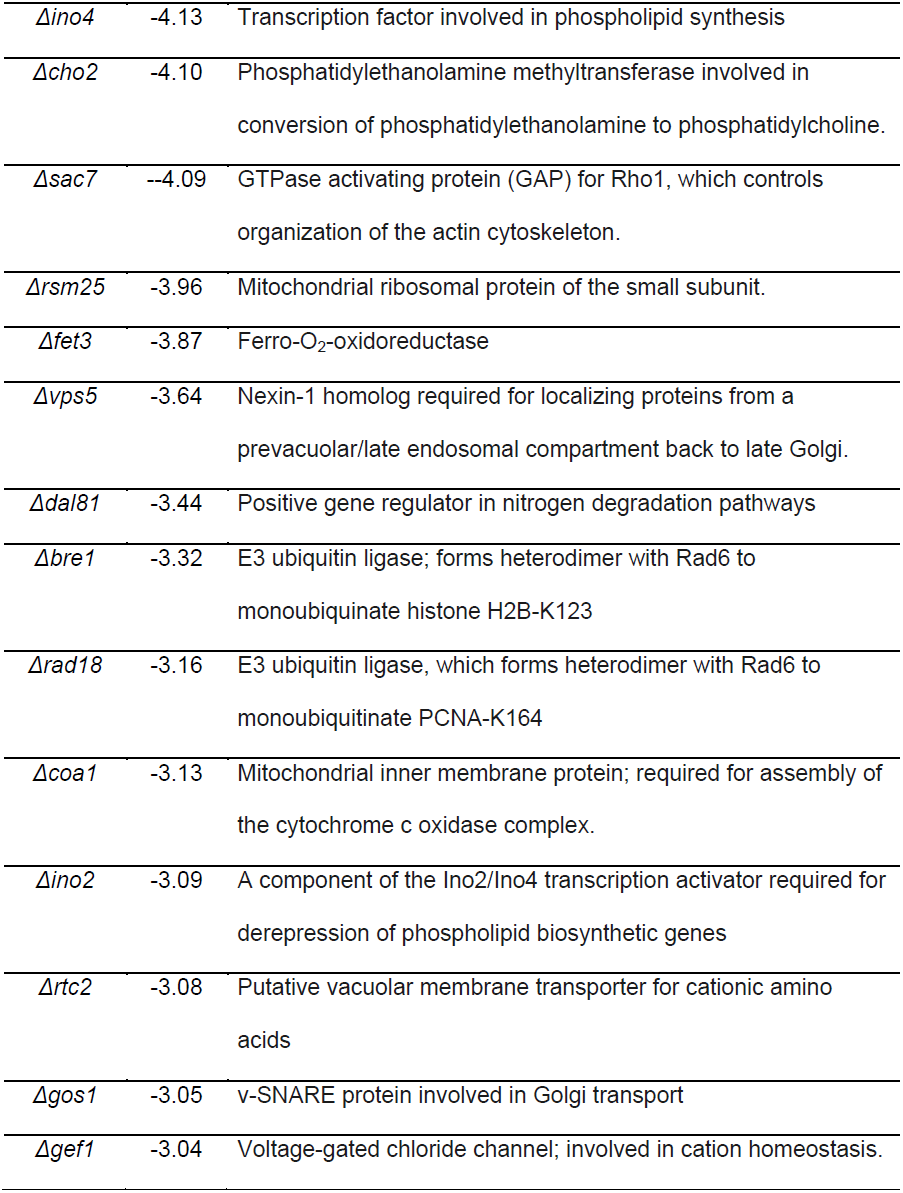
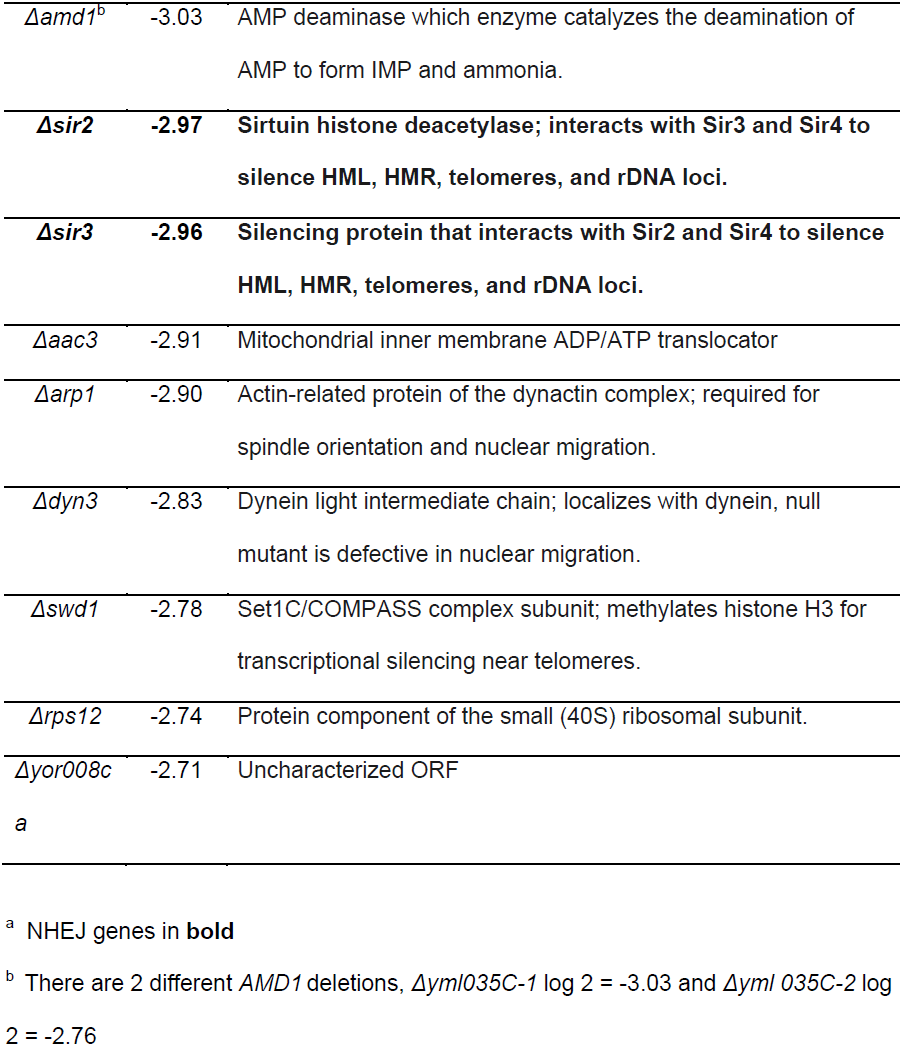
34 genes for which deletion results in a decrease in log2 ratio of cells present after selection for excision compared to cells present after transposase induction decreased log2 < 2.7.

### Construction of pWS2-Integration

pWSY2-Integration is a pRS414-based plasmid containing the *piggyBat* transposase gene under the control of the GALS promoter, mini-*piggyBat-Nat* and *URA3* in the plasmid backbone. The plasmid containing GALS-*piggyBat* transposase and *URA3* generated in the construction of pWS1-Excision was modified by the introduction of mini-*piggyBat-Nat* into the plasmid backbone by homologous recombination, using a PCR fragment containing mini-*piggyBat*-*Nat* flanked by 45 bp of plasmid sequence, selecting for resistance to clonNAT. To facilitate plasmid loss, a G to C point mutation was introduced by site-directed mutagenesis at position 8 of the *CDEI* element of the plasmid *CEN6* site.

### Construction of pWS2-Integration Δtransposase

The *piggyBat* transposase gene was removed by digestion of pWS2-Integration with BamHI and XhoI which cut at the 5’- and 3’-termini of the gene, respectively, followed by fill-in with T4 DNA polymerase and ligation.

### *piggyBat* integration assay

BY4727 pWSY2-Integration which contains mini-*piggyBat-Nat* or BY4727 pWS2-Integration Δtransposase colonies were isolated by growth on SC + 2% galactose + 50 µg/ml clonNAT plates for 5 days and then resuspended, serially diluted and plated on SC + 2% glucose + 1 mg/ml 5-FOA plates to determine the total number of cells and on SC + 2% glucose + 1 mg/ml 5-FOA + 50 µg/ml clonNAT plates, to determine the number of cells in which excision followed by integration occurred. The integration frequency is cells with an integration/total cells.

## RESULTS

### Development of a one-plasmid assay system for *piggyBat* excision

To assay *piggyBat* excision and donor site repair in yeast, we constructed pWSY1-Excision, a pRS414-based plasmid containing the *piggyBat* transposase gene under the control of the GALS promoter and a mini-*piggyBat* transposon, which contains a NatMX resistance cassette, inserted within a yeast *URA3* gene, forming *ura3::* mini-*piggyBat-Nat*. Because *URA3* is disrupted by the mini-*piggyBat-Nat* transposon, a *ura3* auxotrophic strain containing this plasmid remains a uracil auxotroph. *piggyBac* elements such as *piggyBat* excise precisely, restoring the donor site to its pre-transposon sequence (Mitra *et al*. 2008; Mitra *et al*. 2013). Thus when *piggyBat* excises from the *ura3::*mini-*piggyBat-Nat* donor site and the gapped donor site is repaired, *URA3* is expressed and the strain becomes a uracil prototroph (Figure 2). This provides a simple Ura^−^ to Ura^+^ assay for transposon excision and donor site repair following galactose induction of the *piggyBat* transposase in the *ura3Δ0* BY4727 strain.

**Fig 2.**
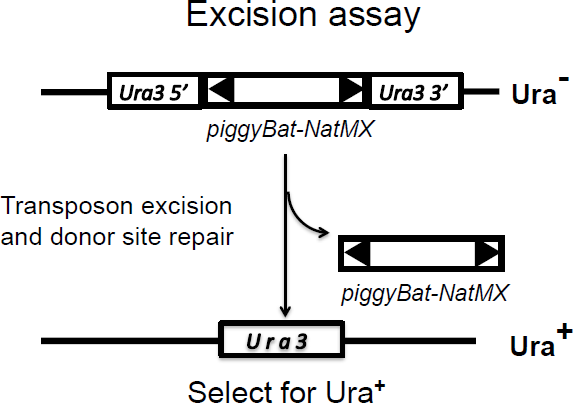
Schematic of the *piggyBat* excision assay. *piggyBat* is excised from *ura3*::mini-*piggyBat-Nat* upon transposase induction and the gapped donor backbone is repaired to generate *URA3*, the cell being converted from a uracil auxotroph to a uracil prototroph.

We observed a mini-*piggyBat-Nat* excision frequency of about 2.9 × 10^−2^ Ura^+^ cells/total cells upon galactose induction of the *piggyBat* transposase gene, a frequency about a thousand-fold higher than the background frequency of 3.5 × 10^−5^ Ura^+^ cells/total cells without transposase induction. The excision frequency of *piggyBat* reported here with a single plasmid system is about 10-fold higher than that previously reported using two-plasmid *piggyBat* excision system (Mitra *et al*. 2013).

### A genome-wide screen suggests that *piggyBat* excision and donor site repair is decreased in strains lacking components of the NHEJ pathway

To identify host factors involved in *piggyBat* transposition, we screened a pool of yeast haploid non-essential single gene deletion strains (Giaever *et al*. 2002) transformed with pWS1-Excision for mutant strains in which transposon excision and/or donor site repair was altered. To identify deletion strains in which transposition was altered, we compared the fraction of each deletion in the transformant pool following transposase induction to the fraction of each deletion strain in the pool following selection for transposon excision and donor site repair. Genes whose deletion result in a decrease in their fraction in the deletion pool after transposition likely encode host factors required for excision and/or donor site repair. By contrast, genes whose deletion result in an increase in their fraction in the deletion pool likely encode host factors that inhibit transposition.

We performed three independent biological replicates starting from transformation of the haploid deletion pool with pWS1-Excision, selecting for plasmid-based clonNAT resistance on SC + 2% glucose + 50 µg/ml clonNAT plates and obtained at least 3 million transformants for each replicate, i.e. about 600-fold coverage of the ∼ 5000 single gene deletion library. Following recovery of the transformants in liquid, expression of transposase was induced by growth in SC + 2% galactose + 50 µg/ml clonNAT. To select cells in which *piggyBat* excision and repair of the *ura3::*mini-*piggyBat-Nat* donor site to *URA3* occurred, the induced transformants were grown for several cycles in SC - Ura + 2% glucose.

We determined the abundance of each deletion strain in the transformant pools in which transposase was expressed by growth in galactose (SC + 2% galactose) and the pools in which *piggyBat* excision and donor site repair occurred by growth in absence of uracil (SC – Ura + 2% glucose). For each pool, we recovered the deletion library barcodes by PCR and quantitatively compared the strains in these pools by next generation sequencing.

Analysis of the barcodes in the three replicate pools following transposase induction revealed that 4,043 of the mutants had at least one read per pool and ∼ 3,300 mutant strains had at least 5 reads per pool. Thus plasmid transformation and growth in SC + 2% glucose + 50 µg/ml clonNAT, followed by transposase induction in SC + 2% galactose, resulted in the loss of about 20% of the original deletion mutant pool.

We also determined the number of reads for each deletion strain in the three replicate pools grown in the absence of uracil (SC - Ura + 2% glucose) to select for transposon excisants. For each replicate, we calculated the log2 ratio of the number of reads present before and after selection for *piggyBat* excision for each strain. The 4043 deletion strains present after selection for excision and their log2 ratios are shown in Supplemental Table S1. Comparison of the log2 ratios of the three replicates gave correlation coefficients of about 0.6 (Supplemental Figure S1) and thus we averaged the log2 ratios from the three replicates.

34 of the 4,043 deletion strains had log2 ratios after selection for excision compared to those after transposase induction that decreased > 2.7 fold (Table 1). These genes are candidates for genes that encode host factors required for transposition. No gene candidate for host factors that inhibit transposition was identified as no deletion strain had a log2 ratio > 2.0 (Supplemental Table S1).

Notably, upon selection for Ura^+^ excisants, the log2 ratio of *Δura2* and *Δura4* strains decreased ∼ 6.17-fold and 4.31-fold, respectively (Table 1). Both these genes are required for uracil biosynthesis (Benoist *et al*. 2000; Lacroute 1968). When assayed in individual *Δura2* and *Δura4* deletion strains (Figure 3), *piggyBat* excision assayed by Ura^+^ selection decreased more than 10,000-fold. Thus our selection for transposase-dependent Ura^+^ excisants successfully identified deletion mutants that blocked formation of Ura^+^ products.

**Fig 3.**
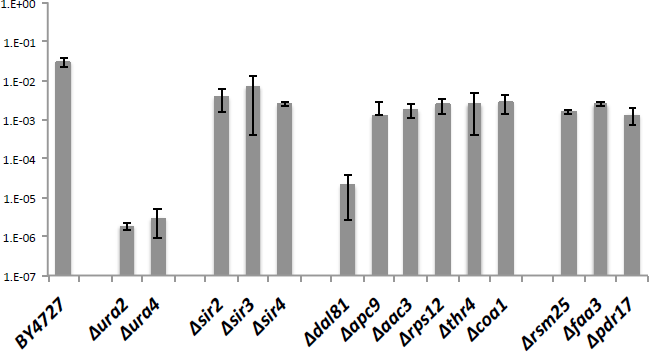
The *piggyBat* excision frequency assayed in individual deletion mutants. *piggyBat* excision was determined using pWS1-Excision in the indicated deletion mutant strains. The average frequency of three independent experiments is shown.

Omitting the *Δura2* and *Δura4* mutants, and also *Δynl296W*, *Δydl041W and Δynr005C because* their ORFs are dubious, we used FunSpec (http://funspec.med.utoronto.ca) to explore the cellular functions of the genes deleted in the strains whose log2 ratio decreased > 2.7-fold upon selection for excision. We found that genes for the GO Biological Process Category Double-strand break repair via non-homologous end joining (NHEJ) [GO:0006303] were significantly enriched (P = 8.6e-12) among these 32 genes. Our screen identified 7 of 25 genes in this category, including *DNL4*, *LIF1*, *NEJ1*, *YKU70*, *YKU80, SIR2*, and *SIR3* (Table 1). As described below, *piggyBat* excision and donor site repair were defective in strains individually deleted for NHEJ genes.

Why weren’t the other 17 genes involved in NHEJ-mediated repair identified in our screen? Recall that only 4,043 of the initial library of 5171 deletion strains were present in the library after growth in SC + 2% galactose. Notably, strains deleted for the NHEJ genes *FYV6*, *MRE11*, *POL2*, *RSC2*, *SIN3* and*VPS75* were not present among these 4043 strains (Supplemental Table 1). Although strains deleted for the NHEJ *genes DOA1*, *IRC20*, *MCK1*, *POL4*, *RAD27*, *RAD50*, *RSC1, RTT109*, *SIR4, SRS2, SUB1* and *XRS2* were present among these 4043 deletion strains, the decreases in their log2 ratios in the induced compared to the excisant pools were not > 2.7 (Supplemental Table 1).

We also assayed *piggyBat* excision and donor site repair in deletion strains individually deleted for the 25 non-NHEJ and non-Ura genes identified in Table 1 except for *Δrtc2, Δswd1 and Δyor008c-a*. Excision was not significantly decreased in most of these strains (Supplemental Figure S2). However, excision was significantly (≥ 10-fold) reduced in six of these 22 mutants, *Δdal81*, *Δapc9*, *Δaac3, Δrps12, Δthr4* and *Δcoa1* (Figure 3). We also observed that excision was significantly decreased (≥ 10-fold) in the deletion strains *Δrsm25, Δfaa3 and Δpdr17* in the course of other experiments. Why deletion of these genes affects *piggyBat* excision and donor site repair remains to be determined.

### *piggyBat* excision is defective in individual strains lacking components of the NHEJ pathway

The core components of the NHEJ repair machinery (Dudasova *et al*. 2004) identified in our screen are the DNA end binding protein Ku complex (*YKU70*, *YKU80*), the DNA ligase IV (*DNL4)* and its associated proteins Lif1/Xrcc4 (*LIF1*), and Nej1 (*NEJ1)*, which is the yeast ortholog of the mammalian NHEJ component XLF (XRCC4-like factor; Cernunnos). To verify that these NHEJ pathway genes are indeed involved in donor site repair after *piggyBat* excision, we measured the excision frequency of *piggyBat* in individual *Δyku70*, *Δyku80, Δdnl4, Δlif1* and *Δnej1* strains using the pWSY1-Excision assay, finding that the frequency of *piggyBat* excision in the *Δyku70*, *Δyku80 Δdnl4*, and *Δlif1* strains was about 10,000-fold lower than that in the parental BY4727 strain. Excision in the *Δnej1* strain is about 100-fold lower than that in the parental BY4727 (Figure 4).

**Fig 4.**
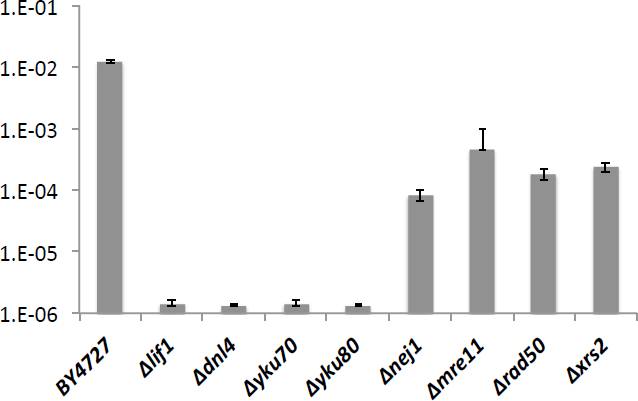
The *piggyBat* excision frequency assayed in individual NHEJ deletion mutants. *piggyBat* excision was determined using pWS1-Excision in the indicated NHEJ deletion mutant strains. The average frequency of three independent experiments is shown.

NHEJ in yeast also requires Mre11, Rad50 and Xrs2, which form the MRX complex (Rine and Herskowitz 1987; Trujillo *et al*. 2003). However, strains with these deletions were not present in the pre-excision deletion pool after growth in SC + 2% galactose + 50 µg/ml clonNat (Supplemental Table 1). Analysis of *piggyBat* excision in strains individually deleted for these genes revealed that excision was about 100-fold lower in these strains than in the parental BY4727 strain (Figure 4).

### *piggyBat* excision and donor site repair is reduced in diploids

The NHEJ pathway is down-regulated in diploids (Valencia *et al*. 2001). Although the starting single gene deletion strains in our screen were haploids, diploid formation can occur when mating type switching is de-repressed. Such de-repression, diploid formation and hence reduction of NHEJ repair, occurs in strains defective for Sir2, Sir3, and Sir4, which repress the mating type loci (Rine and Herskowitz 1987). We observed in our screen that *piggyBat* excision was decreased in *Δsir2*, and Δ*sir3* strains (Table 1). *piggyBat* excision in individual *Δsir2*, *Δsir3*, and *Δsir4* strains, was decreased about 10-fold compared to excision in the parental haploid strain (Figure 3).

We also examined *piggyBat* excision and donor site repair in a diploid generated by mating the haploid BY4727 *MATa* pWSY1-Excision strain with the haploid BY4741 *MATα* strain, finding that *piggyBat* excision and donor site repair in the diploid strain was about 20-fold lower than that in a haploid strain (Table 2).

**Table 2.** *piggyBat* excision frequency in haploid and diploid cells.

| Strain | Without | With |
| --- | --- | --- |
|  | transposase induction | transposase induction |
| Haploid | $1.30 \pm 0.09 \times 10^{-4}$ | $2.11 \pm 0.08 \times 10^{-2}$ |
| Diploid | $1.86 \pm 2.54 \times 10^{-5}$ | $1.26 \pm 0.53 \times 10^{-3}$ |

*piggyBat* excision frequency was determined using pWS1-Excision

Thus the results of our screen and assays in individual deletion strains implicate the NHEJ pathway in gapped donor site repair after *piggyBat* excision. Although it might be argued that from the results described thus far, the NHEJ pathway is required for *piggyBat* excision *per se* rather than donor site repair, we show below that *piggyBat* does excise in NHEJ mutants.

### NHEJ repair is not required for *piggyBat* integration

The excised *piggyBat* transposon has 3’OHs on its 3’ ends and TTAA extensions on its 5’ ends (Figure 1). During integration the 3’OH ends attack both strands of the target DNA at staggered positions, resulting in covalent attachment of the 3’ *piggyBat* ends to the target DNA. One pathway for the regeneration of intact duplex DNA at the insertion site would be for the TTAAs on the 5’ transposon ends to anneal to the complementary TTAA sequences of the target DNA, followed by ligation. One potential source of this ligation activity is NHEJ Dnl IV ligase complex. Thus we analyzed *piggyBat* integration in NHEJ mutants.

To assay integration, we constructed pWSY2-Integration, a pRS414-based plasmid containing the *piggyBat* transposase gene under the control of the GALS promoter, a mini-*piggyBat-Nat* transposon, and *URA3* in the plasmid backbone. Integration was measured by selecting for cells that have lost the plasmid by growth on 5-fluoroorotic acid (5-FOA^R^) and in which the mini-*piggyBat*-*Nat* had jumped to the chromosome (clonNat^R^) (Figure 5). The observed *piggyBat* integration frequency in BY4727 is about 5.8 × 10^−2^ 5-FOA^R^ clonNat^R^ cells/ total cells (Figure 6). We found that the frequency of *piggyBat* integration in all these NHEJ *Δyku70*, *Δyku8, Δdnl4, Δlif1*, and *Δnej1* mutant strains was about the same as that in parental strain BY4727. Therefore we conclude that the ligation machinery provided by the NHEJ pathway is not required for the repair of *piggyBac* integrants.

**Fig 5.**
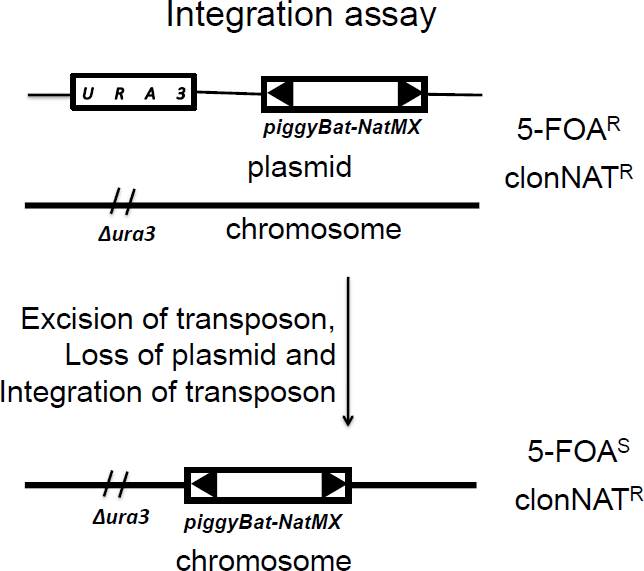
The *piggyBat* integration assay. Mini-*piggyBat-Nat* on a *URA3* plasmid is excised upon transposase induction and can integrate in the chromosome. Integration events are assayed as cells that retain the mini-*piggyBat-Nat* (Clon^R^) but have lost the donor plasmid (5-FOA^R^).

**Fig 6.**
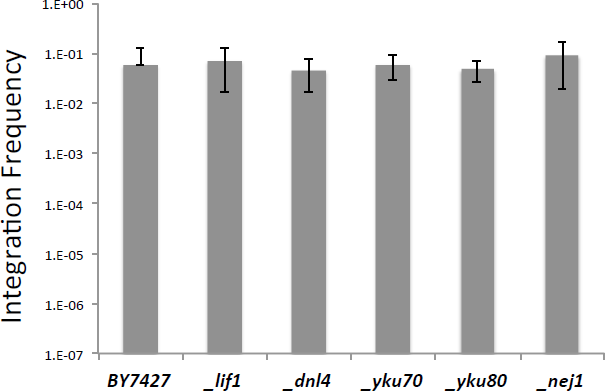
*piggyBat* integration frequency in individual NHEJ deletion mutants. *piggyBat* integration using pWS2-Integration was measured in the indicated deletion strains. The average integration frequency of three independent experiments is shown. *piggyBat* excision frequency was also assayed in each deletion strain using pWS2-Integration Δtransposase and the frequency of excision was at least 100-fold lower than with transposase.

The other source of yeast ligase activity is DNA ligase 1 encoded by *CDC9*, which is an essential gene, making its role in *piggyBat* integration difficult to establish (Willer *et al*. 1999).

## Discussion

Here we have explored the requirements for *S. cerevisiae* host factors in transposition of the eukaryotic mammalian cut & paste transposable element *piggyBat*, a member of the widespread *piggyBac* superfamily (Fraser *et al*. 1996; Sarkar *et al*. 2003), which we have previously demonstrated is active in yeast (Mitra *et al*. 2008). We used a single plasmid-based excision reporter system in a high-throughout screen using the haploid yeast single gene deletion collection. We had anticipated that deletion of genes involved in a variety of cellular processes including various DNA-based transactions might affect transposition but found that our screen highlighted only genes involved in NHEJ as being necessary for *piggyBat* excision and donor site repair. A limited dependence on host factors is, however, consistent with the fact that only *piggyBac* transposase is required for transposition *in vitro* (Mitra *et al*. 2008) and that *piggyBac* transposition is highly efficient in a wide variety of organisms ranging from yeast, protozooa, planeria, insects, plants, and a variety of vertebrates including mammals, as well as various types of stem cell (Ding *et al*. 2005; Gonzalez-Estevez *et al*. 2003; Handler 2002; Nishizawa-Yokoi *et al*. 2015). It should be noted, however, that our screen tested involvement of only about 4043 of the haploid non-essential 5171 yeast genes in *piggyBac* transposition, likely because we used minimal synthetic complete media.

The requirement for NHEJ to repair the broken donor backbone from which *piggyBat* has excised is compatible with the known biology and chemical steps of *piggyBac* excision (Mitra *et al*. 2008). *piggyBac* inserts into TTAA sites such that upon *piggyBac* excision, the flanking donor DNA has TTAA overhangs on both 5’ ends of the broken donor DNA (Figure 1). Annealing of these fully complementary TTAA overhangs followed by ligation can restore the donor site to its original TTAA sequence, consistent with *piggyBac* precise excision. Other work (Daley *et al*. 2005) has established that NHEJ in yeast requires the end binding complex MRX (Mre11-Rad50-Xrs2) (Dudasova *et al*. 2004), which is recruited early to DSBs. The core complexe Ku (yKu70-yKu80) binds to and protects the broken ends and the DNL IV ligase (DnlIV-Lif1-Nej1), is recruited by Ku. We have demonstrated that *piggyBat* excision and donor site repair is highly defective in mutant strains deleted for the genes encoding these proteins.

The core NHEJ Ku and DNL IV complex genes are conserved in all eukaryotes (Symington and Gautier 2011) and thus likely mediates repair of the donor site for *piggyBac* transposons in its varied hosts including *piggyBat* in its mammalian host the little brown bat (Mitra *et al*. 2013; Ray *et al*. 2008). Notably, programmed gene assembly in *Paramecium* that is mediated by a domesticated *piggyBac*-like transposase also requires NHEJ (Dubois *et al*. 2012).

The NHEJ pathway is also involved in donor site repair for other DNA cut & paste transposons such as *Ac/Ds* as studied in maize (Rinehart *et al*. 1997) and yeast (Yu *et al*. 2004), and *Sleeping Beauty* in mammalian cells (Walisko *et al*. 2006; Yant and Kay 2003), although in these cases excision does not regenerate the pre-transposon donor site, rather usually leaving footprints that derive from the target site duplications (Rinehart *et al*. 1997; Woodard *et al*. 2012).

The structure of a newly inserted *piggyBac* element is related to the structure of the gapped donor site in that both contain complementary TTAA sequences (Figure 1). The newly inserted transposon has TTAA extensions at its 5’ ends that can anneal to the complementary target DNA TTAAs derived from the staggered joining positions of the transposon to the TTAA target site. Upon TTAA annealing, the newly inserted transposon is flanked by nicks such that formation of intact duplex DNA requires only ligation. Note also that upon *piggyBat* insertion there are only nicks in the new *piggyBat* insertion site that could be sealed by ligation to generate intact duplex DNA. Notably, however, we have found that the NHEJ Dnl IV ligation complex is not required following transposon insertion to generate intact duplex DNA at the insertion site nor are other NHEJ components required. Thus we imagine that the other cellular ligase, DNA ligase 1 which is the *CDC9* gene product, can seal the nicks at the target site. More extensive target site repair than simple ligation is required upon integration of non-target site-specific transposons and retroviral elements because in these cases the 3’ OH target DNA ends are flanked by single strand gaps (Craig 2002) rather than simple nicks as discussed above for *piggyBac* elements. Repair of these flanking gaps could reasonably result from gap-filling by a repair polymerase followed by ligation (Syvanen *et al*. 1982) or possibly by a more elaborate gap-filling mechanism related to that used by bacteriophage *Mu* (Choi and Harshey 2010). We cannot exclude the possibility that repair of a *piggyBac* insertion site may also proceed by removal of the TTAAs at the 5’ transposon ends, and repair of the flanking gaps by a repair polymerase and ligase. Insertion site DNA repair of some bacterial elements has been shown to also require host-mediated disassembly of highly stable post-transposition transposase-DNA complexes (Abdelhakim et al. 2010; North and Nakai 2005). It will be interesting to see if such processes might also be required for DNA repair at eukaryotic transposon insertions sites.

## Acknowledgements

We are grateful to Jef Boeke, Pam Meluh, Kuang Zheng and Sarah Wheelan for their considerable assistance in this project. We also thank Henry Levin for his thoughtful comments on the manuscript and Patti Kodeck for her assistance in preparing the manuscript. NLC is an investigator of the Howard Hughes Medical Institute.

## SUPPLEMENTAL DATA

**Supplemental Figure S1.**
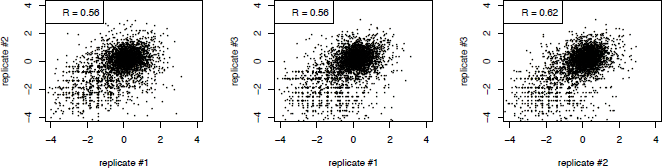
Correlation values for comparison of log2 ratios of barcode sequencing reads from three independent experimental replicates

**Supplemental Figure S2.**
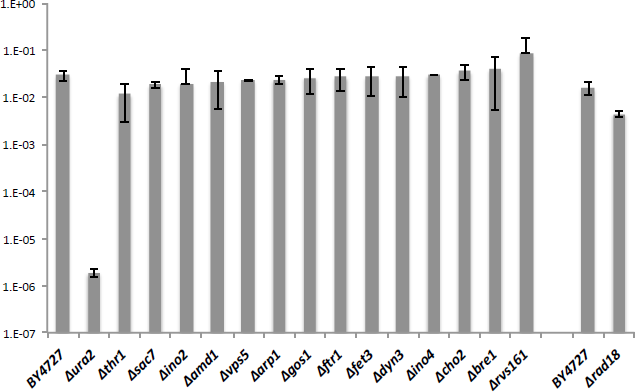
The *piggyBat* excision frequency re-assayed in indicated individual deletion mutants. *piggyBat* excision was determined using pWS1-Excision in the indicated deletion mutant strains. The average frequency of three independent experiments is shown. *Δrad18* was assayed separately from other mutants.

**Supplemental Table S1.** Comparison of the number of gene copies present after selection for excision compared to gene copies present after transposase induction as determined by UPTAG sequencing in three independent experiments

**Supplemental Table 2.**
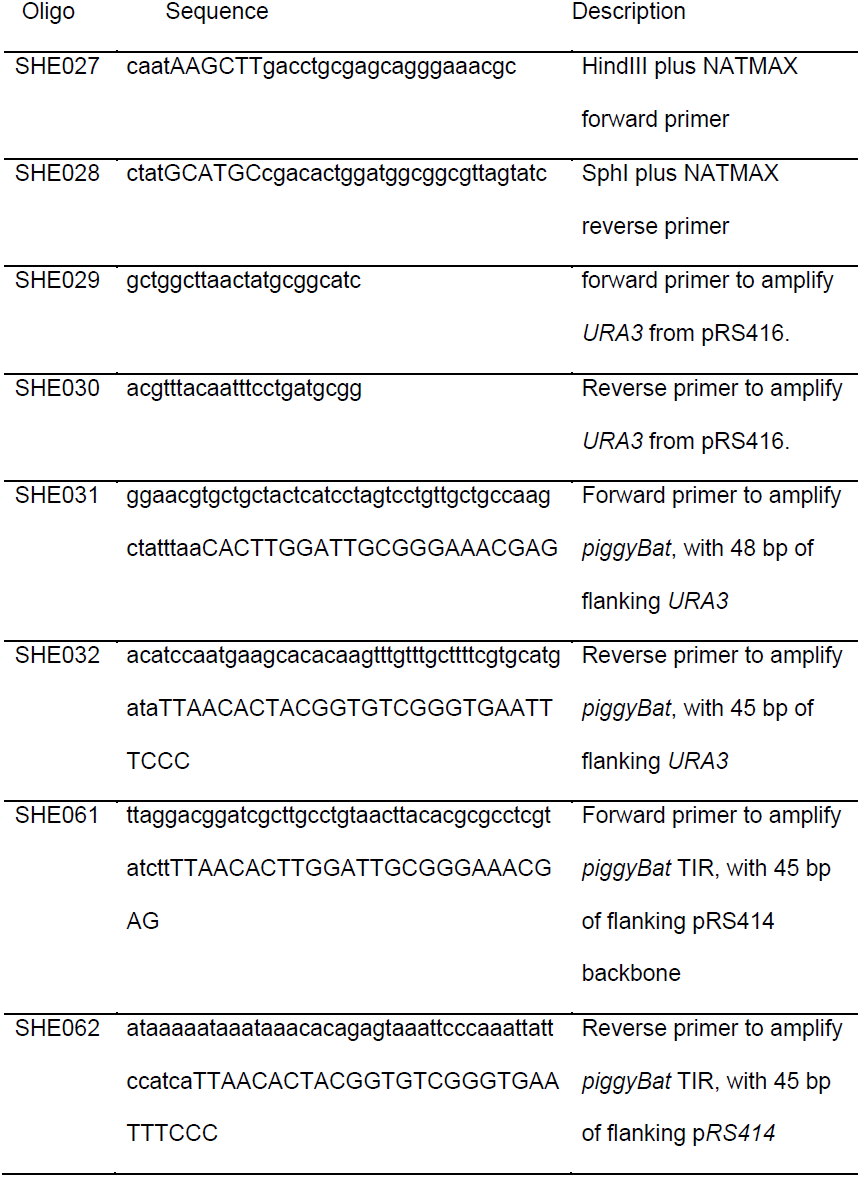
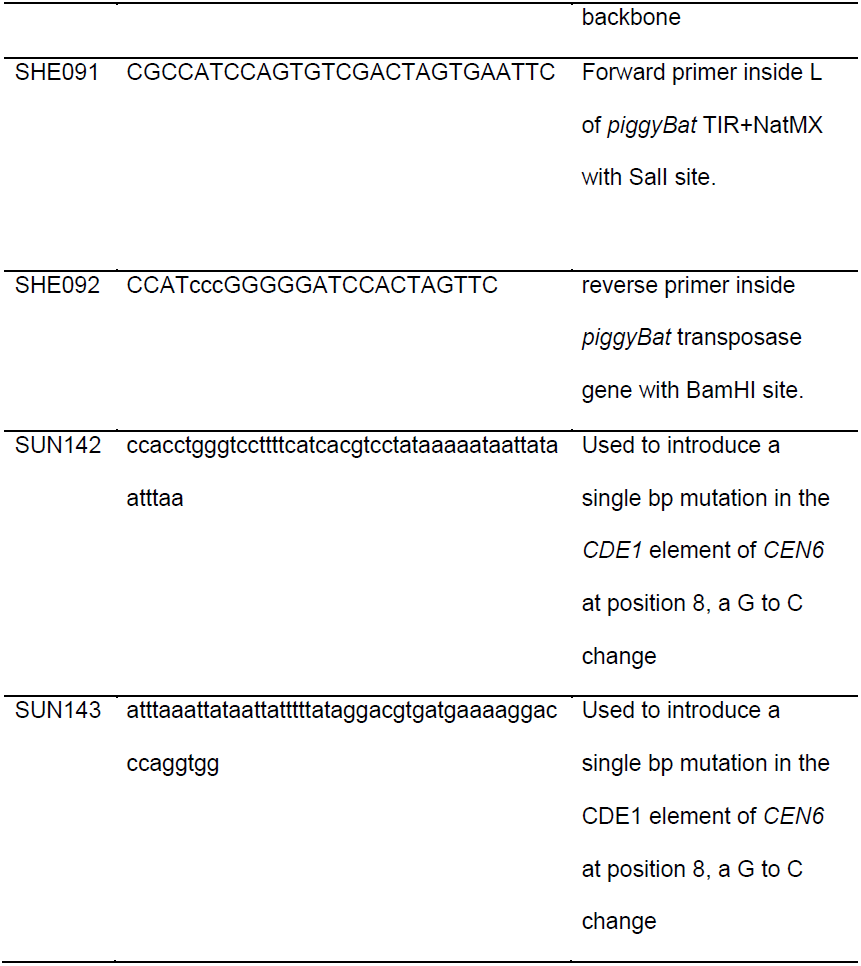
Oligonucleotides

